# Genomic Insights into a Multispecies Bacterial Pathogen Complex Driving Bacterial Blotch in White Button Mushrooms

**DOI:** 10.64898/2026.01.12.699132

**Authors:** Sameerika D Mudiyanselage, Michelle Lee, Jose C. Huguet-Tapia, Romina Gazis, Samuel J Martins

## Abstract

Bacterial blotch remains a major constraint to global white button mushroom (*Agaricus bisporus*) industry, yet its etiological complexity has been underestimated. Through a genome resolved, polyphasic approach applied to symptomatic mushrooms collected from United States, we uncovered an unexpectedly diverse complex of *Pseudomonas* species driving blotch disease. Beyond classical pathogens (*P. tolaasii*, *P. gingeri*, *P. agarici*, and *P.* “*reactans*”, *P. yamanorum*, *Pseudomonas* sp. NC02), our analyses revealed a striking prevalence of *P. azotoformans*, a species not previously associated with mushroom pathology, alongside *P. pergaminensis*, *P. monsensis*, *P. tensinigenes*, *P. simiae*, *Pseudomonas* sp. Irchel 3A7, *Pseudomonas* sp. REP124 and two putatively novel lineages. Comparative genomics demonstrated pronounced heterogeneity in accessory genome content, with *P. azotoformans* exhibiting exceptional genomic plasticity indicative of broad ecological adaptability. Secondary metabolite profiling and white line assays further delineated species-specific chemotaxonomic signatures, underscoring the multifactorial nature of virulence. Collectively, this study provides the most comprehensive genomic and phenotypic characterization of blotch-associated *Pseudomonas* in Northern America, overturning the long-held paradigm of a single dominant pathogen. By establishing that bacterial blotch is multispecies disease complex, our findings redefine its epidemiology and lay the foundation for improved diagnostics strategies in mushroom production systems. The emergence and high prevalence of *P. azotoformans* underscore the limitations of diagnostic protocols focused exclusively on classical blotch pathogens and highlight the need for broader, genomics informed detection strategies. Collectively, this work offers actionable insights to strengthen production resilience and support the sustainability of white button mushroom cultivation as the world’s most economically important specialty food crop.

## 1. INTRODUCTION

The white button mushroom (*A. bisporus*) is one of the fastest growing and most widely cultivated mushroom species globally, with major production hubs in China, the United States, the Netherlands, Poland, Italy, and India. In the United States alone, white button mushrooms represent over 90% of total mushroom consumption underscoring their central role in the national and global mushroom markets (Royse et al., 2017). Beyond its economic importance, *A. bisporus* is a nutritionally dense functional food, providing high-quality proteins, carbohydrates, essential vitamins, minerals and dietary fibers (Atila et al., 2017). It also produces a diverse array of bioactive compounds, including polysaccharides, phenolics, flavonoids, and terpenoids, which exhibit antioxidant, anti-inflammatory, anti-aging, immunomodulatory properties. These compounds have been linked to numerous health benefits, including the reduction of chronic disease risk and cancer prevention (Zhang et al., 2024; Li et al., 2025). Consequently, these combined nutritional and medicinal attributes establish *A. bisporus* as both a staple crop and a critical source of therapeutic compounds. Despite these benefits the commercial production of *A. bisporus* faces a significant and persistent threat from “bacterial blotch”, one of the most destructive postharvest diseases affecting *A. bisporus,* which has impacted commercial production worldwide for decades (Osdaghi et al., 2019; Soler-Rivas et al., 1999).

Bacterial blotch is primarily associated with *Pseudomonas* species, metabolically versatile and ubiquitous bacteria that naturally inhabit the casing layer. Some *Pseudomonas* strains, such as *P. putida* (Eger, 1972) and *P. poae* n12 (Fermor et al., 2000), play beneficial roles in mushroom cultivation promoting *A. bisporus* growth and contributing to primordia formation. However, pathogenic *Pseudomonas* species often persist in casing materials even after pasteurization, leading to disease outbreaks. The bacterial blotch disease is characterized by rapid discoloration of the mushroom cap, pitting, tissue degradation, and, ultimately, extensive postharvest losses. Different blotch phenotypes correspond to different pathogens: *P. tolaasii* causes brown blotch, producing sunken greenish-brown lesions that expand and coalesce (Soler-Rivas et al. 1999), while *P. gingeri* causes ginger blotch, marked by golden-yellow discoloration (Wong and Fletcher, 1979). Beyond these well-known species, few surveys have revealed a much broader diversity of blotch-causing bacteria, including *P. agarici* (Fatmi et al., 2008, Geels et al., 1994; Young, 1970), *P*. “*reactans*” (Wells et al., 1996), *Pseudomonas* sp. NZ17P (Godfrey et al., 2001)*, P costantinii* (Munch et al., 2000; Taparia et al., 2020), *P. yamanorum*, *P. edaphica*, *P. salomonii*, *Pseudomonas* sp. NC02 (Taparia et al. 2020), and *P. extremorientalis* (Huang et al. 2026). A few non-*Pseudomonas* species have been reported in the years to cause blotch-like symptoms, which include *Mycetocola* sp. (Hamidizade et al. 2020), *Serratia liquefaciens*, *S. proteamaculans*, *Pantoea* sp. (Taparia et al. 2020), and *Cedecea neteri* (Huang et al. 2024), highlighting the increasing complexity of this disease.

Pathogenic *Pseudomonas* species employ multiple virulence mechanisms, including extracellular proteases and lipases (Morihara and Homma, 1985), antifungal metabolites (Henkels et al. 2014), and cyclic lipopeptides (CLPs) that enhance motility, surface colonization, competition, and infection. In addition to pathogen-derived factors, the development of blotch symptoms is influenced by the mushroom’s physiological responses, particularly enhanced tyrosinase activity and increased phenolic content, which collectively accelerate browning and tissue discoloration (Soler-Rivas et al. 1999).

Historically, the identification of bacterial blotch-causing *Pseudomonas* species has relied heavily on phenotypic assays such as the White Line Agar Assay (WLA), which detects the interaction between tolaasin produced by *P. tolaasii* and the WLIP compound produced by the organism described as *P. reactans*. However, the increasing diversity of blotch-causing *Pseudomonas* species and the frequent inconsistency of WLA results have made it clear that phenotypic tests alone are insufficient for accurate species identification. Although multilocus sequence analysis (MLSA) and 16S rRNA-based approaches have been widely used for *Pseudomonas* taxonomy, several studies have demonstrated their limitations, particularly in resolving closely related taxa (Gomila et al. 2015). Consequently, modern polyphasic strategies that combine MLSA, whole-genome comparisons, and chemotaxonomic profiling now provide the most robust framework for species-level resolution within the genus *Pseudomonas*.

Despite decades of research, the complete and accurate diversity of blotch-causing *Pseudomonas* species remains unresolved, highlighting the urgent need for broader surveys that address both identification and management, as effective control strategies depend on precise pathogen recognition. This growing complexity underscores the importance of region-specific assessments of pathogen diversity. In this study, we conducted a systematic survey of symptomatic white button mushrooms collected from two commercial production sites in the United States. Specifically, we aimed to characterize the causative agents of bacterial blotch by integrating phenotypic assays (WLA and fluorescence), pathogenicity tests, MLSA, whole-genome analyses, comparative genomics, and *in silico* profiling of secondary metabolite biosynthetic gene clusters. Together, these approaches provided a comprehensive overview of the bacterial blotch pathogens present in these production systems and their associated dynamics on *Agaricus bisporus*.

## 2. MATERIALS AND METHODS

### 2.1. Isolation of *Pseudomonas* sp. from white button mushrooms

White button mushrooms displaying bacterial blotch symptoms were collected between April and May 2025 from two major commercial mushroom production companies in the United States. These companies represent large-scale, vertically integrated production systems operating multiple geographically distributed facilities. For this study, samples were collected from one farm located in the southern United States from each supplier, hereafter referred to as Farm X and Farm Y.

At Farm X, sampling was conducted during the second and third production flushes, whereas at Farm Y, sampling occurred during the first flush. At the time of collection, disease incidence was substantially higher at Farm X (approximately 30%) compared to Farm Y (1–2%). Symptomatic mushrooms were transported to the laboratory under cold conditions in individual ziplock bags, and bacterial isolation was conducted within 24 to 48 h of arrival. Upon arrival, the mushrooms were gently rinsed with sterile distilled water to remove surface debris. Approximately 1 cm² of tissue was excised from the margin between diseased and healthy areas using a sterile scalpel. Each tissue sample was transferred into a microcentrifuge tube containing 1 mL of 10% sodium hypochlorite (NaOCl), vortexed for 2 s, and allowed to stand at room temperature for 30 s for surface disinfection. The samples were subsequently rinsed twice with 1 mL of sterile distilled water, briefly vortexing after each rinse. The disinfected tissue was then macerated in 100 µL of 0.01 M sterile phosphate buffer (pH 7.0) using a sterile disposable pestle, and the homogenate was vortexed to ensure a uniform suspension. Serial dilutions were prepared, and a 10 µL aliquot from each dilution was streaked onto King’s B (KB) (King et al., 1954) agar plates, supplemented with novobiocin (45 µg mL^-1^), penicillin (75 units mL^-1^), and cycloheximide (75 µg mL^-1^) (Suyama et al., 2000). As a control, isolation was also performed from one asymptomatic mushroom collected at each site. All plates were incubated at 27°C for 48 h to allow for bacterial growth. Morphologically distinct bacterial colonies were re-streaked onto freshly prepared KB plates until pure single-colony isolates were obtained. Pure cultures were preserved in 50% glycerol at -80°C for subsequent analyses.

### 2.2. Genomic fingerprinting of bacterial isolates

A total of 76 bacterial isolates were selected for BOX-PCR fingerprinting to assess genetic diversity and identify representative isolates for subsequent analyses. A single colony from each isolate was suspended in 50 µL of sterile distilled water, heated at 95°C for 10 min to lyse the cells, immediately chilled on ice for 2 min, and centrifuged at 12,000 × g for 5 min to pellet cell debris. One microliter of the supernatant was used directly as the DNA template for BOX-PCR amplification using the BOXA1R primer (5′-CTACGGCAAGGCGACGCTGACG-3′), following a modified protocol by Rademaker et al. (1998). The PCR reaction mixture contained 1× Green GoTaq buffer, 0.08 mg mL^-1^ BSA (NEB), 10% DMSO, 1.25 mM dNTPs, 1.76 µM BOXA1R primer, 1.25 U GoTaq G2 DNA polymerase (Promega), and 1 µL of DNA template, for a total volume of 25 µL. PCR amplification was performed in an Applied Biosystems Veriti 96-well thermal cycler with the following cycling conditions: initial denaturation at 95°C for 2 min; 29 cycles of 94°C for 3 s, 92°C for 30 s, 50°C for 1 min, and 65°C for 8 min; followed by a final extension at 65°C for 8 min. Amplified products were separated by electrophoresis on 1.5% (w/v) molecular-grade agarose gels in 0.5× TAE buffer (Quality Biological) at 60 V and 30 mA for 16 h at 4°C. The gels were visualized, and banding patterns were captured using the Biorad gel doc XR imaging system. DNA fingerprint profiles were further analyzed using ImageJ (Sánchez-Jaramillo et al., 2022) for pattern recognition and assessment. Pairwise dissimilarities among isolates were calculated using Jaccard’s distance (Jaccard, 1908), and hierarchical clustering was performed with the unweighted pair group method with arithmetic mean (UPGMA) (Sokal and Sneath, 1963) using R software (v4.4.1). Clusters were defined at a 90% similarity threshold, and a binary heatmap was generated to visualize variation in band distribution among isolates.

### 2.3. Pathogenicity Assay

Based on the genetic dissimilarity analysis, 61 isolates were selected for *in vitro* pathogenicity testing. Bacterial suspensions for inoculation were prepared from 5 mL overnight cultures grown in KB broth at 27°C with shaking at 120 rpm for 24 h. The cultures were centrifuged at 8,000 rpm for 10 min, and the resulting cell pellets were washed once with 0.01 mM phosphate-buffered saline (PBS), followed by centrifugation. The cells were resuspended in the same buffer, and the optical density at 600 nm (OD₆₀₀) was adjusted to 0.5 using a Bio-Rad SmartSpec™ 3000 spectrophotometer, corresponding to approximately 10⁹ colony-forming units (CFU) per mL. *Pseudomonas protegens* BP1304 and *Pseudomonas tolaasii* BP1106 were used as negative and positive controls, respectively. Pathogenicity assays were performed on fresh, healthy white button mushrooms obtained from a local market. To maintain high humidity, sterile plastic trays were lined with moist filter paper. The mushroom caps were surface sterilized by immersion in 10% sodium hypochlorite (NaOCl) for 30 s, rinsed twice with sterile distilled water, and air-dried under sterile conditions. The stalks were removed, and the epidermal layer of the cap surface was peeled away to expose the inner tissue. Caps were then cut into uniform cubes (4 cm × 2 cm) using sterile scalpels, and each cube was placed individually in a sterile petri dish to prevent cross-contamination. Three inoculation sites were designated on each cube using a sterile No. 4 cork borer (4 mm diameter) to mark circular areas, following Henkels et al. (2014). A 10 μL aliquot of bacterial suspension was applied to each marked site. For each isolate, nine replicate mushroom cubes were tested, and the entire assay was independently repeated three times. Petri dishes containing the mushroom cubes were sealed and incubated in the dark at 27°C for three days. Symptom development was monitored and photographed daily to assess tissue maceration, pitting, or discoloration until 72 h. Severity scores were assigned as follows: 1= mild or slow discoloration; 2= rapid tissue discoloration without tissue pitting; 3= extensive discoloration accompanied by tissue pitting.

### 2.4. Florescence and White Line Agar (WLA) test

All isolates (*n* = 61) were cultured on KB medium, and their ability to produce fluorescent pyoverdine and related metabolites was evaluated under UV light (365 nm) after 48 h of incubation. The white line reaction assay was performed to assess the production of extracellular lipopeptides, including tolaasin and WLIP (white line inducing principle), following the methods described by Osdaghi et al. (2019). Briefly, *P. fluorescens* Pf-5P was streaked against each isolate to evaluate tolaasin production, while *Pseudomonas tolaasii* BP1106 was used to assess the secretion of WLIP and related lipopeptides. On King’s B agar, two loopfuls of each test isolate were streaked approximately 8 mm from the reference strain. Plates were incubated at 27 °C, and white-line formation was monitored over a 72 h period. A positive white-line reaction was defined as the formation of a distinct white precipitate at the interface between the test isolate and either *Pseudomonas fluorescens* Pf-5 or *P. tolaasii* BP1106.

### 2.5. DNA extraction and Whole Genome Sequencing

Genomic DNA from the (*n* = 61) isolates was extracted using the Quick-DNA Fecal/Soil Microbe MiniPrep Kit (Zymo Research). The quality and concentration of the DNA were verified using a Bio-Rad Universal Hood Gel Doc XR UV Transilluminator (200 VA) system, and quantity was evaluated using Nanodrop spectrophotometer (Thermo Fisher Scientific). Whole-genome sequencing was performed by SeqCenter (Pittsburgh). Sequencing libraries were constructed using the Illumina DNA Prep Kit (tagmentation-based) with custom IDT 10-bp unique dual indices (UDI) and an average insert size of approximately 280 bp. Paired-end sequencing (2 × 151 bp) was conducted on an Illumina NovaSeq X Plus platform. Raw read demultiplexing, adapter removal, and initial quality control were performed using bcl-convert (v4.2.4).

### 2.6. Genome assembly

The quality of the reads was assessed with FastQC (v0.12.1) (Chen et al., 2018). Based on the quality metrics, low-quality bases and residual adapter sequences were trimmed using Trim Galore (v0.6.10), applying a Phred quality cutoff of 20. To remove potential primer or adapter contamination, 15 bases were clipped from the 5′ ends and 10 bases from the 3′ ends of both forward and reverse reads. Paired reads that passed all quality filters were retained for downstream analyses. High-quality reads were assembled de novo using SPAdes (v4.0.0) (Bankevich et al., 2012), and contigs shorter than 500 bp and a K-mer coverage of less than 2.0 were filtered out. Assembly quality was evaluated with QUAST (v5.2.0) (Gurevich et al., 2013), and genome completeness was estimated using BUSCO (v5.8.3) based on lineage-specific single-copy orthologs (Simão et al., 2015). Only *Pseudomonas* strains (*n* = 57) were used for downstream analysis. Assembled genomes for all 57 strains were submitted to NCBI under Bio Project PRJNA1377199, with accession numbers SAMN53759319 to SAMN53759375.

### 2.7. Genome-wide comparisons

The identity of each isolate was determined using data from assembled genome sequences submitted to the Type (Strain) Genome Server TYGS, (https://tygs.dsmz.de) for whole-genome-based taxonomic analysis(Meier-Kolthoff and Göker, 2019; Meier-Kolthoff et al., 2013). TYGS calculates digital DNA–DNA hybridization (dDDH) values using multiple formulas, including dDDH4, which estimates nucleotide identity within high-scoring genome alignments. A dDDH4 value above 70% is used for taxonomic delineation (Meier-Kolthoff et al., 2013). For further classification, the Average Nucleotide Identity (ANI) based Genome Taxonomy Database classify workflow GTDB-Tk (v2.4.1) (Chaumeil et al., 2022) was used with a threshold of 95% to determine species boundaries. TYGS and GTDB-Tk were also used to identify the closest relative strains, including type strains, for comparative analysis. Pairwise ANI values of the isolates and their closest relatives were computed using all-versus-all strategies with FastANI (v1.34) (Dataset S02) (Jain et al., 2018). ANI similarity values greater than 95% were used to construct a phylogenetic heatmap in RStudio using the pheatmap package (v1.0.13). When the above methods were insufficient to confidently resolve species-level identity, we further assessed genomic relatedness using the Genome-to-Genome Distance Calculator (GGDC) (https://ggdc.dsmz.de/ggdc.php) (Meier-Kolthoff et al., 2022).

Multi-locus sequence analysis (MLSA) was performed to infer the evolutionary history of the bacterial genomes using eight conserved housekeeping genes (*rpoD, rpoB, recA, gapA, gltA, atpD, gyrB,* and *ileS*) from nine reference strains (Dataset S01). A total of 31 closely related reference sequences, including *Pseudomonas lurida* LMG21995^T^ as the outgroup, were retrieved from NCBI and included in the phylogenetic analysis (Dataset S01). Reference strains included all available type strains unless otherwise noted. Type strain names were confirmed with The List of Prokaryotic names with Standing in Nomenclature (LPSN) (Parte et al., 2020). Gene sequences were extracted and aligned using AutoMLSA2 (v2.0.9.0) (https://github.com/davised/automlsa2), which features an automated pipeline that identifies housekeeping genes, performs multiple sequence alignment using MAFFT (v7.520), and concatenates the alignments. The concatenated alignment totaled 176,583 bp and was used to generate a maximum-likelihood phylogenetic tree using IQ-TREE (v3.0.1) with node support values based on 1,000 bootstraps (Wong et al., 2025, Minh et al., 2020). The phylogenetic tree was visualized using iTOL (v7.2.2) (Letunic et al., 2024).

### 2.8. Core and Pan genome analysis

Pan-genome analysis was performed on all 57 *Pseudomonas* isolates, without incorporating external NCBI reference genomes. Additionally, for the four most abundant species *P. tolaasii*, *P. gingeri*, *P. azotoformans*, and *P. pergaminensis* pan-genome and core-genome profiles were constructed by integrating corresponding NCBI reference genomes. All the NCBI reference accessions are listed in Dataset S01. Gene prediction and functional annotation were performed with Prokka v1.14.6 (Seemann, 2014) with default parameters. The resulting GFF files were used for the pan-genome analysis with Roary v3.13.0 (Page et al., 2015), applying a minimum BLASTp identity of 90% for both species cluster analyses and for all strains. For each species group, maximum-likelihood phylogenetic analysis was conducted using RAxML v8.2.10 based on the concatenated core-gene alignments generated by Roary. This analysis employed the GTR+GAMMA substitution model with rapid bootstrap searches, with 100 bootstrap replicates to identify the best-scoring tree (Stamatakis, 2014). Phylogenetic trees were visualized and annotated in iTOL (v7.2.2). Both the resulting tree and gene presence/absence matrices were explored and visualized using Phandango (Hadfield et al., 2018).

### 2.9. Secondary metabolite prediction

Secondary metabolite biosynthetic gene clusters (BGCs) were identified using antiSMASH v7.0.0 (Blin et al., 2023) with default parameters. To focus on conserved clusters, only BGCs with over 30% sequence similarity were retained for further analysis. The resulting BGC profiles were compiled for all isolates, and the presence or absence of each predicted cluster type was visualized in R (v4.4.1) using ggplot2 to compare secondary metabolite cluster distributions among the species groups.

## 3. RESULTS

### 3.1. Isolation of *Pseudomonas* spp. from white button mushrooms

During the mushroom survey, we collected 28 symptomatic white button mushrooms from Farm X and 17 from farm Y, both exhibiting typical bacterial blotch symptoms. The symptoms varied in severity, ranging from light yellow, known as “ginger blotch”, to dark brown or chocolate-colored lesions referred to as “brown blotch”. These symptoms were often accompanied by sunken, water-soaked areas and collapse of the wet tissue (Fig. 1A). A total of 76 bacterial isolates were selectively recovered from the samples using NCP-KB medium and were used for subsequent analyses.

**Fig. 1.**
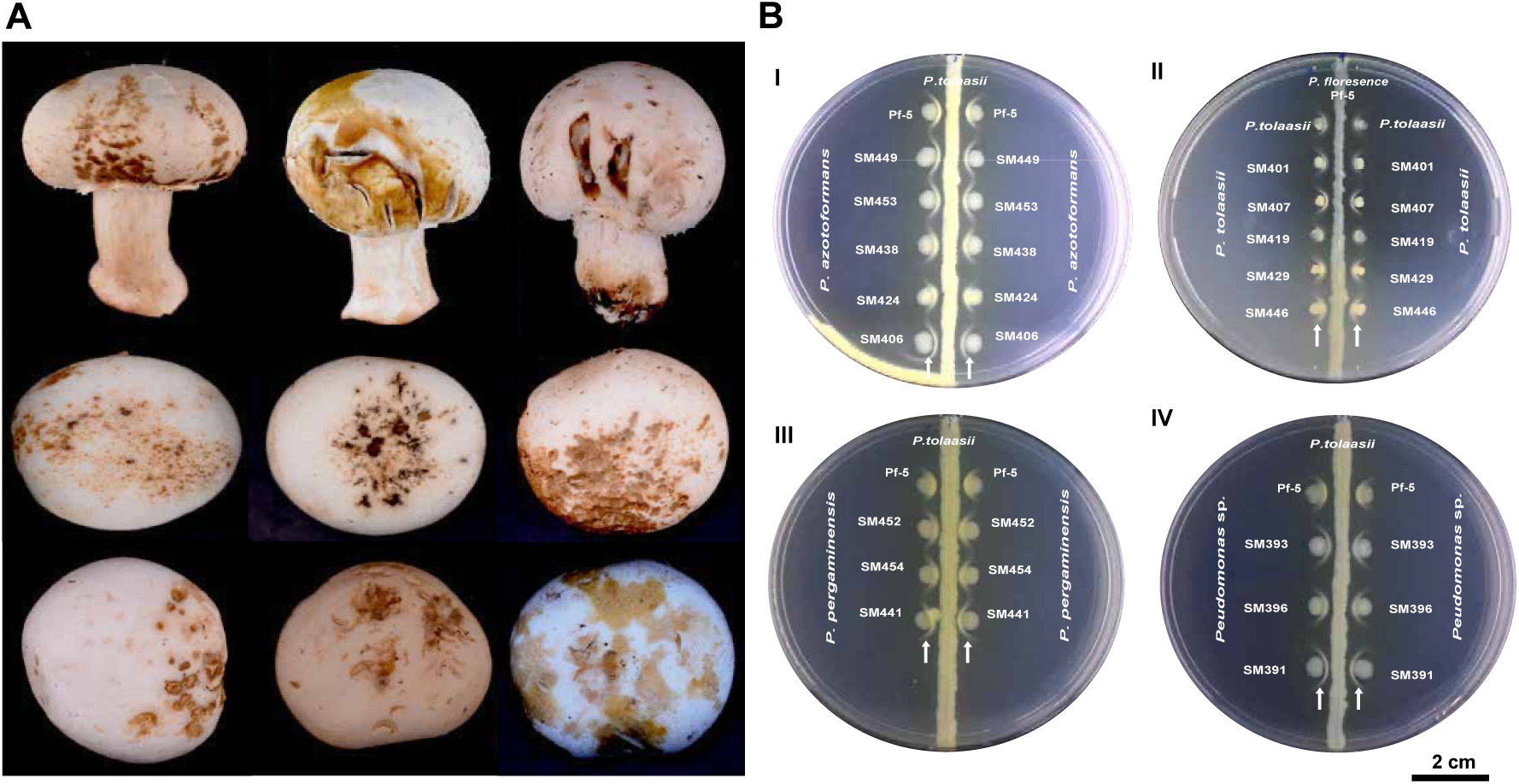
Phenotypic characterization of bacterial blotch pathogens from white button mushroom collected at two U.S. farms (X and Y) in the U.S. (A) Representative blotch symptoms on mushroom caps, ranging from brown to ginger discoloration with varying degrees of tissue pitting. (B) White-line reaction assays illustrating lipopeptide-mediated interactions among *Pseudomonas* isolates: B(I) *P. azotoformans* versus *P. tolaasii* BP1106; B(II) *P. tolaasii* isolates versus *P. fluorescens* Pf-5; B(III) *P. pergaminensis* isolates versus *P. tolaasii* BP1106; B(IV) isolates SM391, SM393, and SM396 versus *P. tolaasii* BP1106. Arrows highlight distinct white precipitates, indicating positive interactions associated with cyclic lipopeptide production. Scale bar = 2 cm.

To identify isogenic strains prior to further genomic and phenotypic characterization, BOX-PCR fingerprinting was performed on the 76 bacterial isolates recovered from the symptomatic mushrooms. The BOX-PCR profiles exhibited banding patterns in agarose gel ranging from <0.5 kb to approximately 10 kb. A binary data matrix representing the presence or absence of 23 distinct BOX-PCR bands was generated and subjected to clustering analysis. The resulting dendrogram (Supplementary Fig. S1) revealed 60 isolate groups at a 90% similarity threshold, representing 76 distinct isolates with varying degrees of relatedness. This clustering pattern demonstrated a high level of genetic diversity among the isolates while also identifying 10 groups of closely related strains (isolate group 8, 14, 18, 23, 26, 37, 39, 40, 57, and 58). Based on these DNA fingerprint patterns, 15 isolates exhibiting identical BOX-PCR profiles were considered isogenic and excluded from subsequent analyses.

### 3.2. Pathogenicity Assay

The ability of each isolate to cause bacterial blotch symptoms was assessed *in vitro* by inoculating fresh mushroom tissue cubes with each of the 61 selected isolates from the BOX-PCR profiles. In this study, pathogenicity was defined as the ability to induce discoloration on mushroom tissue within 2 days post-inoculation, in contrast to the controls. A total of 85% of the strains were identified as producing extensive discoloration accompanied by tissue pitting. An additional 7% of isolates produced only rapid tissue discoloration without tissue pitting, while 8% exhibited only mild or slow discoloration, comparable to the nonpathogenic control strains *Pseudomonas protegens* BP1304 and *Pseudomonas tolaasii* BP1106 within 48 h post-inoculation (Table 1).

**Table 1.**
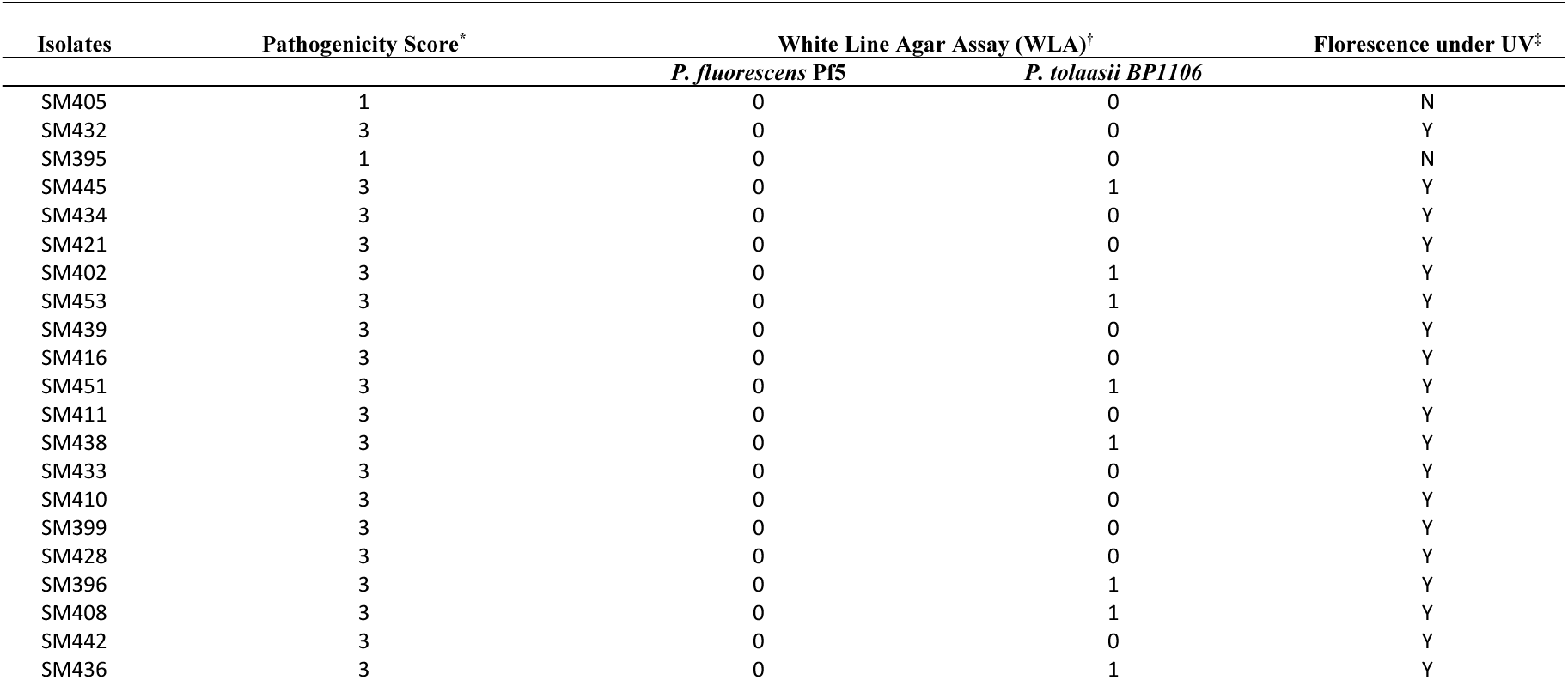
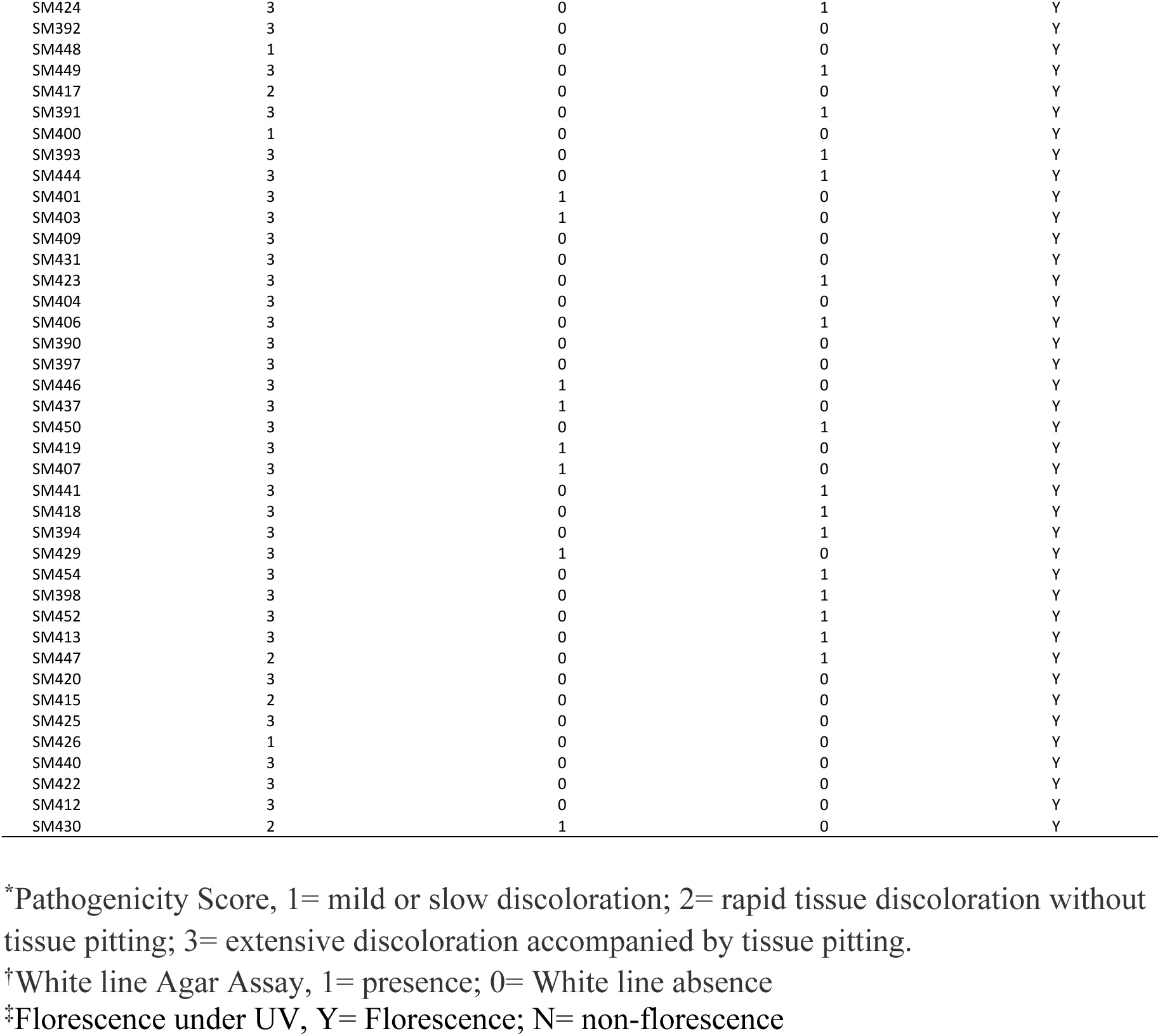
Phenotypic characterization of isolates based on pathogenicity, white-line agar Assay (WLA), and fluorescence production.

| Isolates | Pathogenicity Score* | White Line Agar Assay (WLA)* |  | Fluorescence under UV‡ |
| --- | --- | --- | --- | --- |
|  |  | *P. fluorescens* Pf5 | *P. tolaasii* BP1106 |  |
| SM405 | 1 | 0 | 0 | N |
| SM432 | 3 | 0 | 0 | Y |
| SM395 | 1 | 0 | 0 | N |
| SM445 | 3 | 0 | 1 | Y |
| SM434 | 3 | 0 | 0 | Y |
| SM421 | 3 | 0 | 0 | Y |
| SM402 | 3 | 0 | 1 | Y |
| SM453 | 3 | 0 | 1 | Y |
| SM439 | 3 | 0 | 0 | Y |
| SM416 | 3 | 0 | 0 | Y |
| SM451 | 3 | 0 | 1 | Y |
| SM411 | 3 | 0 | 0 | Y |
| SM438 | 3 | 0 | 1 | Y |
| SM433 | 3 | 0 | 0 | Y |
| SM410 | 3 | 0 | 0 | Y |
| SM399 | 3 | 0 | 0 | Y |
| SM428 | 3 | 0 | 0 | Y |
| SM396 | 3 | 0 | 1 | Y |
| SM408 | 3 | 0 | 1 | Y |
| SM442 | 3 | 0 | 0 | Y |
| SM436 | 3 | 0 | 1 | Y |
| SM424 | 3 | 0 | 1 | Y |
| SM392 | 3 | 0 | 0 | Y |
| SM448 | 1 | 0 | 0 | Y |
| SM449 | 3 | 0 | 1 | Y |
| SM417 | 2 | 0 | 0 | Y |
| SM391 | 3 | 0 | 1 | Y |
| SM400 | 1 | 0 | 0 | Y |
| SM393 | 3 | 0 | 1 | Y |
| SM444 | 3 | 0 | 1 | Y |
| SM401 | 3 | 1 | 0 | Y |
| SM403 | 3 | 1 | 0 | Y |
| SM409 | 3 | 0 | 0 | Y |
| SM431 | 3 | 0 | 0 | Y |
| SM423 | 3 | 0 | 1 | Y |
| SM404 | 3 | 0 | 0 | Y |
| SM406 | 3 | 0 | 1 | Y |
| SM390 | 3 | 0 | 0 | Y |
| SM397 | 3 | 0 | 0 | Y |
| SM446 | 3 | 1 | 0 | Y |
| SM437 | 3 | 1 | 0 | Y |
| SM450 | 3 | 0 | 1 | Y |
| SM419 | 3 | 1 | 0 | Y |
| SM407 | 3 | 1 | 0 | Y |
| SM441 | 3 | 0 | 1 | Y |
| SM418 | 3 | 0 | 1 | Y |
| SM394 | 3 | 0 | 1 | Y |
| SM429 | 3 | 1 | 0 | Y |
| SM454 | 3 | 0 | 1 | Y |
| SM398 | 3 | 0 | 1 | Y |
| SM452 | 3 | 0 | 1 | Y |
| SM413 | 3 | 0 | 1 | Y |
| SM447 | 2 | 0 | 1 | Y |
| SM420 | 3 | 0 | 0 | Y |
| SM415 | 2 | 0 | 0 | Y |
| SM425 | 3 | 0 | 0 | Y |
| SM426 | 1 | 0 | 0 | Y |
| SM440 | 3 | 0 | 0 | Y |
| SM422 | 3 | 0 | 0 | Y |
| SM412 | 3 | 0 | 0 | Y |
| SM430 | 2 | 1 | 0 | Y |

\*Pathogenicity Score, 1= mild or slow discoloration; 2= rapid tissue discoloration without tissue pitting; 3= extensive discoloration accompanied by tissue pitting.
†White line Agar Assay, 1= presence; 0= White line absence
‡Florescence under UV, Y= Florescence; N= non-florescence

### 3.3. Evaluating the potential for lipopeptide and fluorescent pigment production

Among the 61 bacterial isolates examined, eight produced a white-line reaction when confronted with *Pseudomonas fluorescens* Pf-5, indicating the presence of tolaasin-like lipopeptides typically associated with *P. tolaasii* or closely related species. In contrast, twenty-four isolates formed a distinct white line when challenged with *P. tolaasii* BP1106, consistent with the production of WLIP or WLIP-like noncanonical lipopeptides. The remaining 29 isolates showed no reaction with either *P. fluorescens* Pf5 or *P. tolaasii* BP1106 (Table 1, Fig. 1B). Nearly all isolates exhibited fluorescence under UV illumination (365 nm), indicative of pyoverdine production, except for two isolates (SM395 and SM405), which did not fluoresce (Table 1).

### 3.4. Whole genome sequencing of *Pseudomonas* isolates

We analyzed the whole genomes of 57 pathogenic *Pseudomonas* strains isolated from *Agaricus bisporus*. Illumina paired-end sequencing produced an average of 1.77 million read pairs per genome, with a range of 1.0 to 3.4 million. Over 95% of bases exceeded a Phred quality score of Q30. De novo assemblies yielded an average of 185 contigs per genome, with a mean total length of 6.77 Mb, N50 values averaging 96,095 bp, and a GC content of 61%. BUSCO analysis indicated an average genome completeness of 98%, with no detected genome duplication, fragmentation, or missing regions, ensuring high-quality data suitable for genome assembly and downstream analyses. Dataset S02 provides summary statistics of the assembly.

### 3.5. Genome-wide comparison and Taxonomic classification

To ensure accurate taxonomic classification and avoid potential misidentifications associated with conventional methods, we employed a polyphasic approach that integrates several complementary analyses. The *Pseudomonas* isolates were first evaluated using the Genome Taxonomy Database Toolkit (GTDB-Tk) and the Type Strain Genome Server (TYGS) for whole-genome-based taxonomic assignment. Additionally, we performed multilocus sequence analysis (MLSA) of 8 conserved housekeeping genes and FastANI-based average nucleotide identity comparisons to confirm species-level classifications. This integrated approach provided a highly reliable and comprehensive strategy for resolving taxonomic relationships within the *Pseudomonas* genus.

We initially used GTDB-Tk for whole-genome classification, relying on GTDB-Tk-based ANI measurements to achieve high-resolution species-level assignments of the isolates. We identified *Pseudomonas “gingeri*” (*n* = 11), *P. azotoformans* (*n* = 10), *P. tolaasii* (*n* = 7), *P. pergaminensis* (*n* = 6), and *Pseudomonas “reactans”* (*n* = 2), as well as single isolates of *P. agarici, P. simiae, P. monsensis, P. yamanorum*, and *P. tensinigenes*. A species boundary of >95% ANI and <83% for interspecies divergence was adopted. Isolate SM400 shared 94% GTDB-Tk-based ANI with *P. allokribbensis*, while SM426 exhibited 90% ANI with *P. azerbaijanoccidens*, indicating low relatedness. In total, 14 isolates could not be confidently assigned to any *Pseudomonas* species and were not further explored in this study that focused on *Pseudomonas* species. Among these, SM412, SM422, SM425, and SM440 showed >95% ANI to *Pseudomonas* sp. NC02, while SM415 and SM430 showed >95% to *Pseudomonas* sp. Irchel 3A7 and *Pseudomonas* sp. REP124, respectively. The remaining isolates SM390, SM391, SM393, SM396, SM397, SM404, SM409, and SM431 did not show >95% ANI to any species represented in the GTDB-Tk database, indicating that they may represent previously uncharacterized *Pseudomonas* lineages.

We further characterized the isolates using the TYGS, which utilizes a curated database of prokaryotic type strains for whole-genome-based taxonomic classification. This analysis identified isolates belonging to *P. tolaasii, P. pergaminensis, P. agarici, P. simiae, P. monsensis,* and *P. yamanorum*, with digital DNA-DNA hybridization (dDDH) values exceeding the 70% species delineation threshold (Meier-Kolthoff et al. 2013), consistent with the results obtained from GTDB-TK classification. In contrast, the *P. tensinigenes* isolate showed a slightly lower dDDH value (66%). However, due to the absence of corresponding valid type strains in the TYGS database, *Pseudomonas “gingeri”*, *Pseudomonas* “*reactans*”, and *P. azotoformans* could not be validated. Additionally, the remaining 14 *Pseudomonas* isolates could not be resolved to any known type strain in the database. The complete list of ANI and dDDH values for all isolates is provided in Dataset S01.

To assess genetic relatedness among mushroom-associated *Pseudomonas* isolates, we conducted pairwise ANI analyses. Using a ≥95% ANI threshold for species delineation, the dataset resolved into 14 distinct genomic clusters, capturing the full spectrum of intra and interspecific diversity within the collection (Fig. 2A). Comparative analyses with closely related reference type strains enabled the assignment of multiple clusters to recognize species, including *P. “gingeri”, P. azotoformans, P. tolaasii, P. pergaminensis, P. “reactans”, P. agarici, P. simiae, P. monsensis, P. yamanorum*, and *P. tensinigenes*, with clustering patterns aligning with taxonomic classifications derived from TYGS and GTDB analyses. Isolates SM400 and SM426 did not cluster with *P. allokribbensis* and *P. azerbaijanoccidens*, respectively, indicating significant genomic divergence from these reference strains. The remaining 14 isolates formed three distinct genomic clusters: Cluster 1 (SM412, SM422, SM425, SM440), Cluster 2 (SM391, SM393, SM396), and Cluster 3 (SM390, SM397, SM404, SM409, SM431), with SM415 and SM430 representing distinct lineages.

**Fig. 2.**
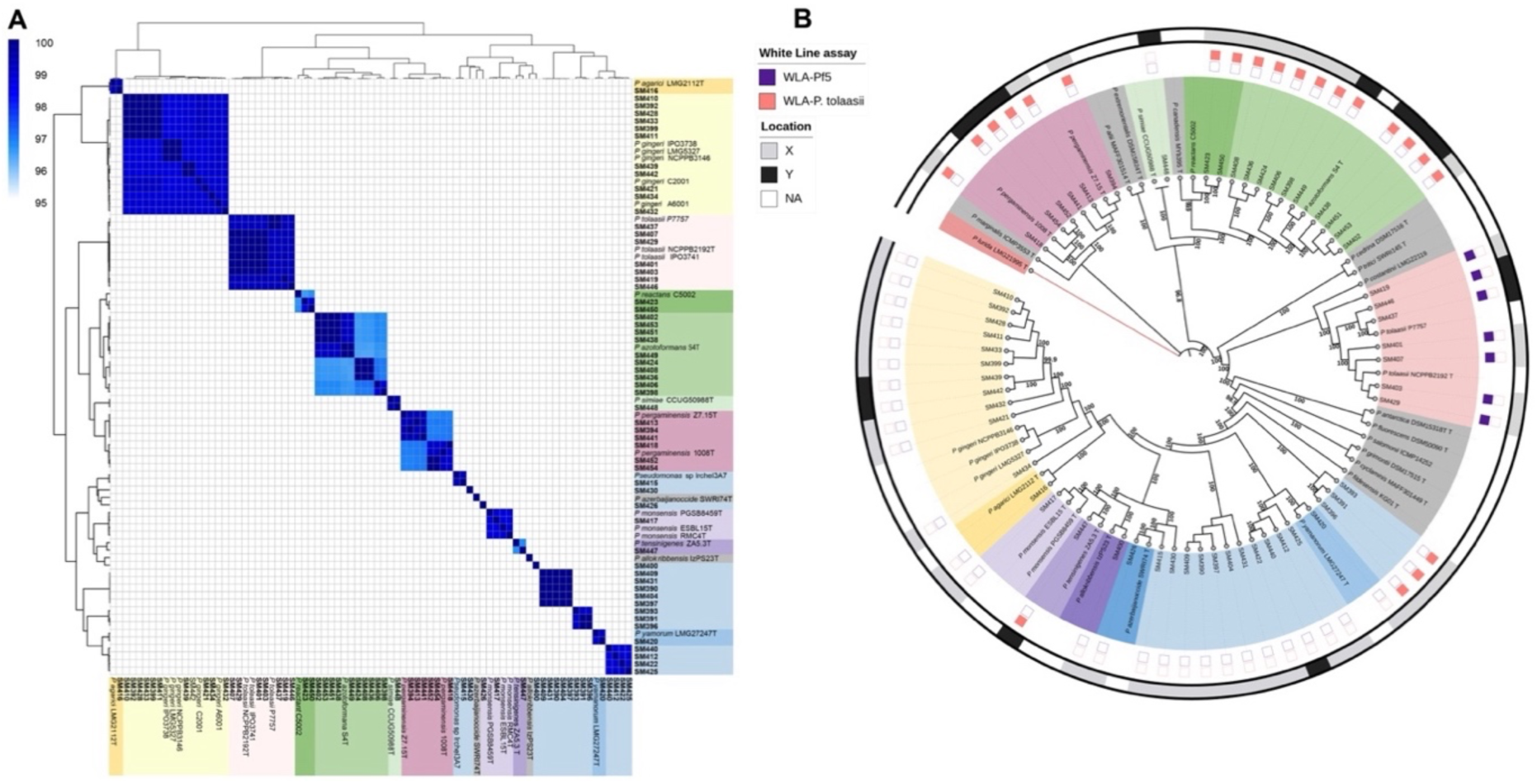
Genomic taxonomy and phylogenetic relationships of *Pseudomonas* isolates associated with bacterial blotch in white button mushrooms (*Agaricus bisporus*) from two U.S. farms (X and Y). (A) Heatmap of pairwise whole-genome Average Nucleotide Identity (ANI) among 57 blotch-associated isolates and 22 reference strains. Color intensity reflects genome similarity, with values ≥95% indicating species-level identity. Axis labels are color-coded by species cluster. (B) Circular maximum-likelihood phylogenetic tree reconstructed from concatenated sequences of eight housekeeping genes (*rpoD, rpoB, recA, gapA, gltA, atpD, gyrB, and ileS*) for 57 isolates and 31 reference strains. Bootstrap support values ≥95% are shown; branch lengths are not to scale*. Pseudomonas lurida* served as the outgroup. The outer ring denotes farm origin (X or Y), while inner markers indicate white-line assay results: pink squares = positive interaction with *P. tolaasii* BP1106; purple squares = positive interaction with *P. fluorescens* Pf*-*5.

### 3.6. Multi-locus sequence analysis

To evaluate the commonly used MLSA for taxonomic resolution and compare it with phylogenomic inference, we selected eight conserved housekeeping genes *rpoD, rpoB, recA, gapA, gltA, atpD, gyrB*, and *ileS* as molecular barcodes based on their established discriminatory power and phylogenetic stability (Gomila et al., 2015). A maximum-likelihood phylogeny was reconstructed from the concatenated sequences of these loci for 57 *Pseudomonas* isolates, including 32 type strains.

The largest lineage corresponded to *Pseudomonas* “*gingeri*”, comprising 11 isolates. Three major sister clades were resolved, including an independent lineage represented by SM434, while the remaining reference strains formed a separate, well-supported subclade. SM416 grouped with the type strain *P. agarici* LMG 2112ᵀ, forming a distinct and clearly separated lineage. Consistent with ANI and GTDB-Tk assignments, seven isolates clustered with *P. tolaasii* NCPPB 2192ᵀ, while six isolates grouped with *P. pergaminensis* 1008ᵀ and Z7.15ᵀ. Similarly, ten isolates clustered with *P. azotoformans* S4ᵀ, and SM423 and SM450 grouped with *P*. “*reactans*” C5002. Several isolates exhibited broader genomic affinities, with *P. monsensis*, *P. tensinigenes*, *P*. *allokribbensis,* and *P. azerbaijanoccida* forming closely related yet distinct branches. Furthermore, SM415 and SM430 resolved into two independent lineages. Additional well-supported isolate clusters included SM412, SM422, SM425, SM440; SM391, SM393, SM396; and SM390, SM397, SM404, SM409, and SM431.

### 3.7. Comparative Pan-Genome Analysis

Roary identified 1,206 core genes and 45,131 accessory genes across the 57 *Pseudomonas* isolates. Among these, 133 soft-core genes were present in at least 95% but fewer than 99% of strains; 9,240 shell genes were present in at least 15% but fewer than 95% of strains; and 35,758 cloud genes were present in at least one strain but fewer than 15%. In total, the pangenome comprised 46,337 genes, highlighting extensive genomic diversity among the isolates. We analyzed the core and pangenome of the four dominant *Pseudomonas* species: *P. tolaasii, P. pergaminensis, P. “gingeri”*, and *P. azotoformans* (Fig. 3). In *P. tolaasii*, the pangenome comprised 8,356 genes, including 5,161 core genes (62%) shared across nearly all isolates, and an accessory genome of 3,195 genes (1,118 shell and 2,077 cloud). *P. pergaminensis* showed a comparable pattern, with a total of 9,092 genes, of which 4,745 (52%) were core; the accessory genome consisted of 2,339 shell and 2,008 cloud genes.

**Fig. 3.**
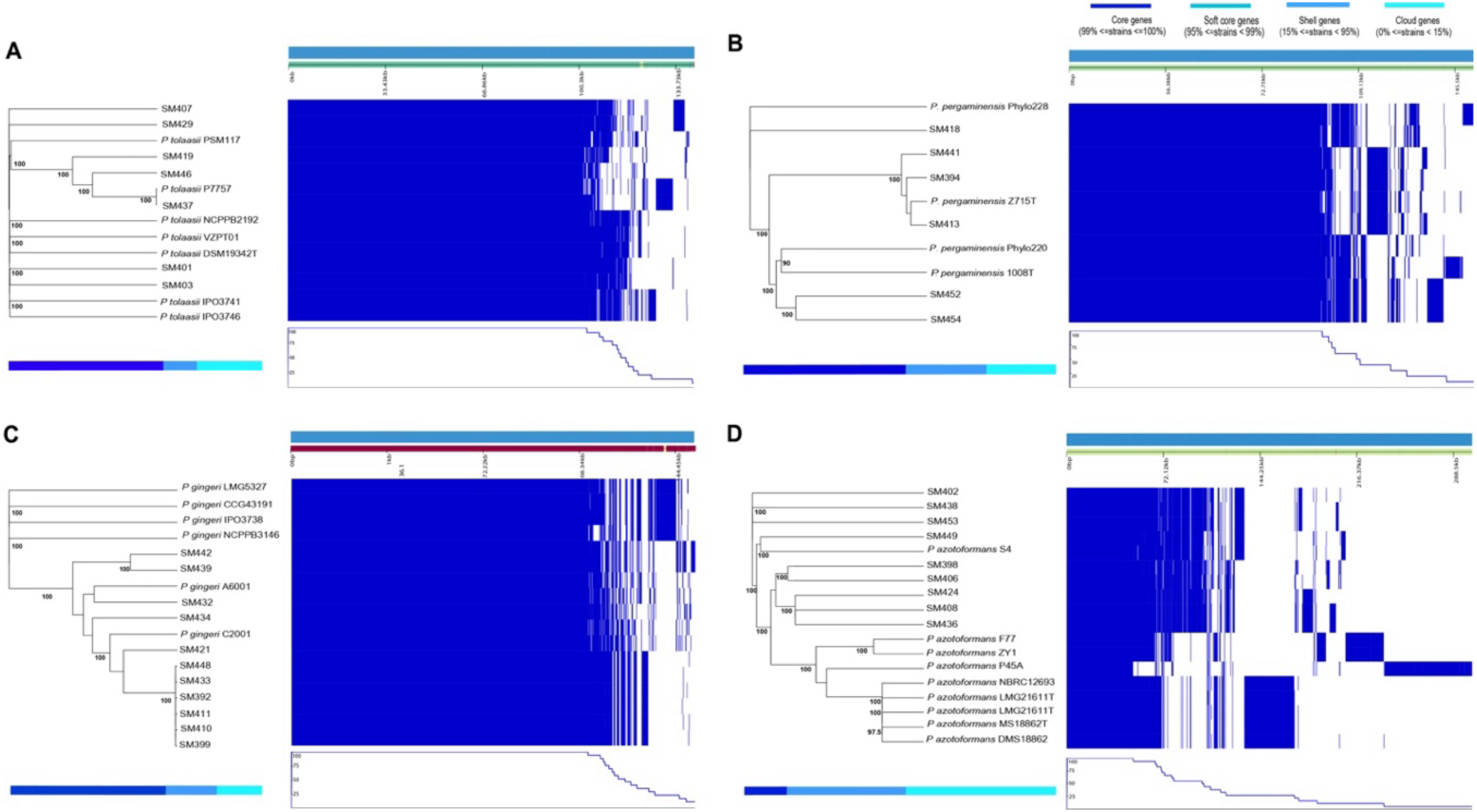
Core and pangenome architecture of four *Pseudomonas* species associated with bacterial blotch in white button mushrooms (*Agaricus bisporus*) from two U.S. farms (X and Y). (A) *P. tolaasii;* (B) *P. pergaminensis;* (C) *Pseudomonas “gingeri”;* (D) *P. azotoformans.* Each panel shows a maximum-likelihood phylogenetic tree based on core gene alignments for the respective species cluster, alongside a gene presence–absence matrix. Rows correspond to individual genomes; columns represent orthologous gene families. Blue blocks indicate gene presence; white blocks indicate absence. Bar graphs summarize the proportion of core, soft-core, shell, and cloud genes, illustrating species-specific genomic diversity and accessory genome expansion.

*P*. “*gingeri*” harbored 9,026 total genes, including 5,560 core genes (62%) and an accessory component of 1,842 shell and 1,624 cloud genes. These species displayed consistently high core-to-pan-genome ratios, indicating that most isolates share a conserved set of essential metabolic and structural functions.

In contrast, *P. azotoformans* exhibited a markedly expansive pan-genome, with 18,029 genes but only 2,477 core genes (14%). Its accessory genome was dominated by 6,858 shell and 8,694 cloud genes, reflecting extensive genomic variability and a substantially more flexible gene repertoire.

### 3.8. Diversity and distribution of biosynthetic gene clusters among *Pseudomonas* isolates

Beyond DNA-based taxonomy, we generated *in silico* secondary metabolite profiles to examine chemotaxonomic patterns among blotch-associated *Pseudomonas* species. Genome mining of the 57 *Pseudomonas* isolates using antiSMASH revealed a substantial biosynthetic diversity, encompassing 22 distinct families of secondary metabolite biosynthetic gene clusters (BGC) (Fig. 4). The presence-absence matrix illustrates the distribution of major BGC classes, including numerous nonribosomal peptide synthetase (NRPS) loci encoding cyclic lipopeptides such as tolaasin, viscosin, ririwpeptide, lokisin, syringafactin, putisolvin, orfamides B, syringocin, histicorrugatin, necroxime, stechlisin, sessilin, bananamide, and corpeptin-like families. Siderophore-associated clusters were abundant and phylogenetically widespread, including EDHA, yersiniabactin, Ni-siderophore, enantio-pyochelin, and azotobactin D. Several isolates also encoded BGCs for volatile or antibiotic metabolites such as sodorifen, geosmin, and 2,4-diacetylphloroglucinol. Distinct chemotaxonomic signatures emerged across species: *P. gingeri* encoded volatile and antibiotic loci; *P. pergaminensis* consistently carried viscosin; *P. tolaasii* harbored tolaasin; and viscosin clusters were widely distributed across several lineages, including *P. “reactans”*. Cluster-specific patterns were evident, with Cluster 1 enriched in enantio-pyochelin and Cluster 3 encoding combinations of tolaasin and riripeptide like loci. Notably, *P. azotoformans* lacked known BGCs for CLPs, siderophores, volatiles, or antibiotics, suggesting unexplored or cryptic metabolic pathways.

**Fig. 4.**
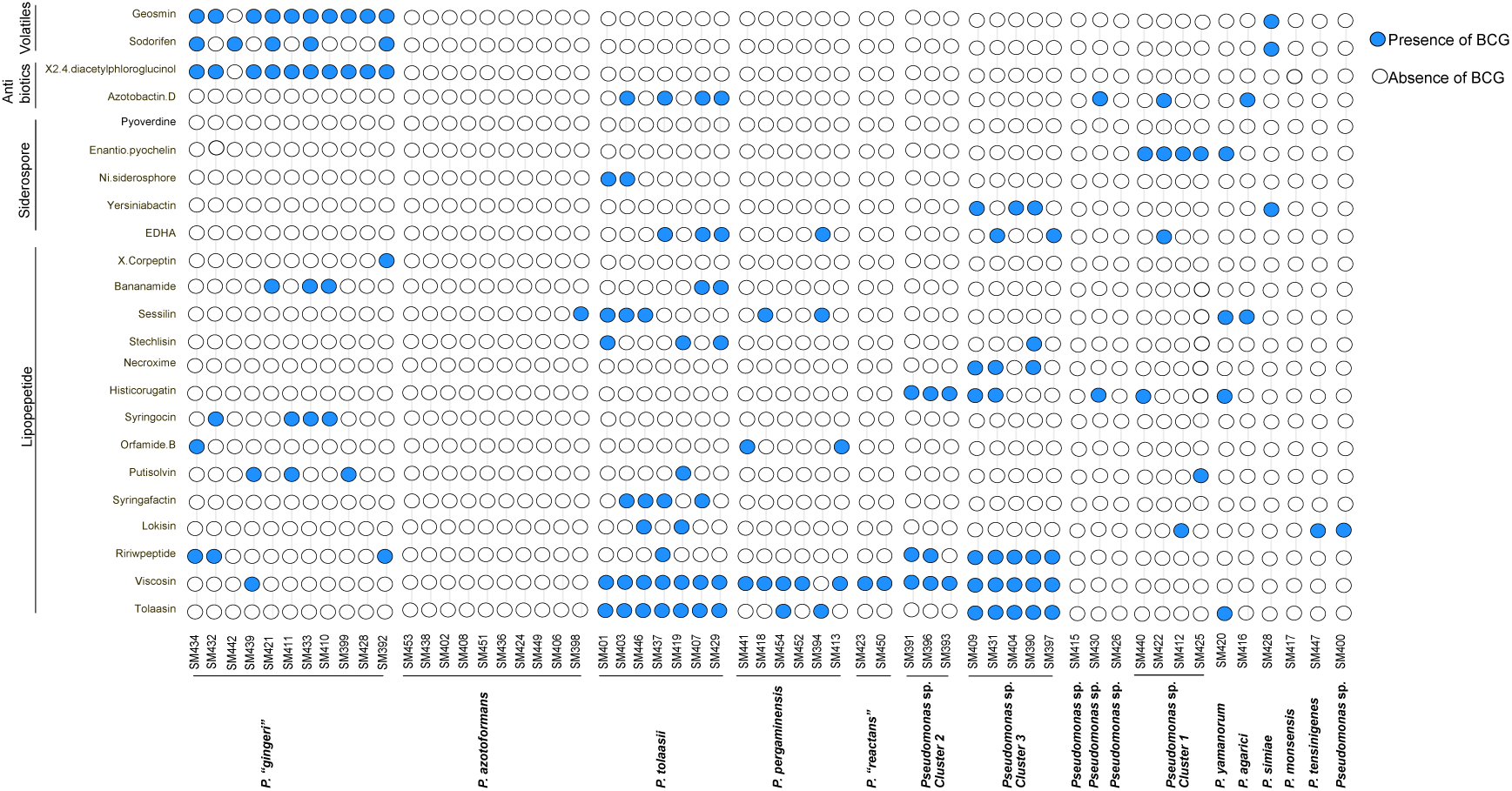
Distribution of 22 Biosynthetic gene cluster (BGC) families across 57 *Pseudomonas* isolates. Columns represent individual isolates, and rows correspond to BGCs families grouped into four functional categories: lipopeptides, siderophores, antibiotics, and volatiles. Blue dots indicate predicted presence of a BGC family with ≥30% sequence similarity as predicted by antiSMASH, while white dots indicate absence. Presence may include both complete and partial clusters, as predicted by antiSMASH.

### 3.9. Pathogenicity assessment

To assess the range of symptom patterns we monitored disease progression over a 48-hour period after inoculation. In Fig. 5 we summarized the symptoms induced by individual *Pseudomonas* species on mushroom cap tissues. Apart from the characteristic “brown blotch” and “ginger blotch” symptoms caused by *P. tolaasii* and *Pseudomonas* “gingeri,” respectively, the additional species identified in this study produced distinct lesion phenotypes. For instance, *P. pergaminensis* induced lighter brown lesions accompanied by pronounced tissue pitting, whereas *P. yamanorum* generated pale yellow discoloration, also associated with noticeable pitting. In contrast, isolates belonging to Cluster 3 were mainly associated with dark, chocolate-brown discoloration and fine, needle-like pitting as the infection progressed. Isolates from Cluster 1 produced pale brown lesions with limited pitting but evident tissue damage. *P. azotoformans* caused irregular brown blotches with slight pitting progression, while *P. agarici* produced wet, yellowish tissue with little or no pitting. *P. monsensis* and *P. tensinigenes* both exhibited dark black discoloration without significant tissue damage or pitting. Furthermore, *P*. *simiae,* isolates in Cluster 2, SM400, and SM426 caused only slight discoloration compared to the nonpathogenic control.

**Fig. 5.**
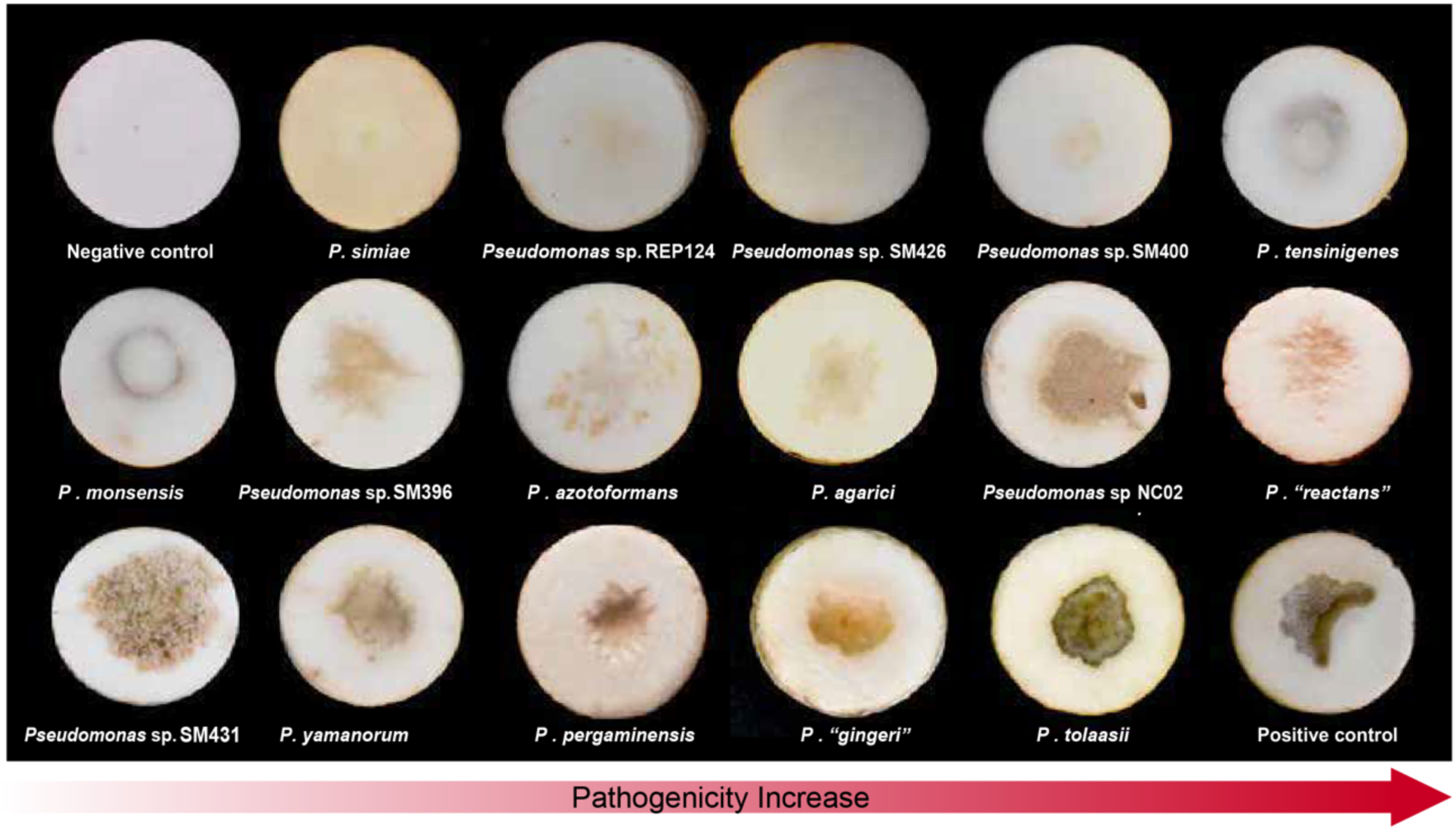
Pathogenicity assays of representative *Pseudomonas* species clusters inoculated on fresh white button mushroom tissue. Lesion phenotypes range from extensive discoloration with pronounced tissue pitting to rapid surface browning and mild or slow discoloration. Arrows indicate increasing pathogenic severity across species clusters.

## 4. DISCUSSION

In this study, we isolated and characterized bacterial blotch-causing *Pseudomonas* from white button mushrooms collected from two southern states in the USA. Among the 56 isolates, we identified well-known blotch pathogens, including *Pseudomonas “gingeri*” (*n* = 11), *P. tolaasii* (*n* = 7), *Pseudomonas* sp. NC02 (*n* = 4), and *Pseudomonas “reactans”* (*n* = 2), along with single isolates of *P. yamanorum*, and *P. agarici*. Additionally, we identified seven bacterial taxa that had not been previously reported as associated with blotch disease, including *P. azotoformans* (*n* = 10), *P. pergaminensis* (*n* = 6)*, P. monsensis*, *P. tensinigenes*, *P. simiae*, *Pseudomonas* sp. Irchel 3A7, and *Pseudomonas* sp. REP124A. Furthermore, we identified two distinct lineages of previously unrecognized *Pseudomonas* spp., representing putative novel species associated with blotch symptoms. The strains SM391, SM393, and SM396 formed one group, while the strains SM390, SM397, SM404, SM409, and SM431 formed another, each constituting well-supported, genomically coherent clades according to ANI, dDDH and MLSA.

Accurate species identification is essential for understanding pathogen diversity and informing disease management strategies. To resolve the taxonomy of the *Pseudomonas* isolates recovered in this study, we applied multiple complementary approaches, including TYGS, GTDB-TK, ANI, and MLSA. Together, these methods provided a robust framework for evaluating genomic relatedness and enabled high-confidence species assignments. Using this integrative strategy, we clearly resolved several blotch-associated taxa, including *Pseudomonas* “gingeri”, *P. tolaasii*, *P.* “*reactans*”, *P. yamanorum*, *P. agarici*, *P. azotoformans*, *P. pergaminensis*, *P. monsensis*, and *P. simiae*, effectively distinguishing our isolates from their closest type strains. However, some groups required more nuanced interpretation. For example, MLSA clustered isolates SM400 and SM426 near *P. allokribbensis* IzPS23ᵀ and *P. azerbaijanoccidentalis* SWRI74ᵀ, yet ANI and GTDB-TK-based dDDH values demonstrated clear genomic distinctiveness, supporting their classification as independent species. Likewise, isolate SM447 showed <70% dDDH similarity to *P. tensinigenes* based on TYGS; however, ANI-based inference provided improved resolution at the species level. A subset of four isolates (SM412, SM422, SM425, SM440) was identified as *Pseudomonas* sp. NC02 according to GTDB-TK. Due to insufficient genome quality for these reference taxa, they could not be fully evaluated using FastANI or MLSA methods. Therefore, we applied GGDC, which confirmed that all four isolates belong to the *Pseudomonas* sp. NC02 lineage. Similarly, SM430 and SM415 identified to be *Pseudomonas* sp. REP124 and *Pseudomonas* sp. Irchel 3A7, respectively. Together, these results highlighted the necessity of multi-method taxonomic frameworks when describing *Pseudomonas* diversity in the mushroom environment. Further, we highlight the inherent differences in resolution between marker gene-based and whole-genome approaches. MLSA, while useful for initial phylogenetic placement and continuity with historical taxonomy, may lack sufficient discriminatory power for closely related taxa due to its reliance on a limited number of conserved loci (Gomila et al., 2015). In contrast, ANI and digital dDDH, which incorporate genome-wide information, provide more robust and reproducible criteria for species delineation. Therefore, in cases of conflict, genome-based approaches should be prioritized, with ANI (≥95%) and dDDH (≥70%) serving as primary criteria for species delineation, while MLSA should be used as a complementary tool to provide phylogenetic context rather than define taxonomic boundaries. BOX-PCR fingerprinting proved valuable as an initial screening tool to capture genomic diversity across all 76 blotch-associated isolates. Although BOX-PCR revealed substantial heterogeneity, its clustering patterns did not correspond with ANI or MLSA-based phylogenies, reflecting differences in repetitive element dynamics rather than true evolutionary relationships. As seen in other bacterial genera such as *Streptomyces*, BOX-PCR is well-suited for rapid delineation of non-redundant isolates but offers limited resolution for deeper taxonomic or phylogenetic inference (Nayak et al., 2011; Rademaker et al., 1998; Van Belkum et al.,1996). Accordingly, in this study, it served primarily as a high-throughput preselection method prior to whole-genome analyses.

Among the novel blotch-associated taxa identified in this study, *Pseudomonas azotoformans* was particularly abundant, representing a notable expansion of the known ecological range of this species. *P. azotoformans* has primarily described in plant-associated contexts, most notably as a pathogen of cereal crops such as rice (Iizuka and Komagata 1963), and as an antagonist of several important plant pathogens, including *Fusarium fujikuroi* (Wiemann et al., 2013), *Colletotrichum* spp. (Popescu et al., 2024), *Pseudomonas syringae* pv. *actinidiae* (Correia et al., 2025), and *Alternaria brassicae* (Bhattacharya et al., 2025). This host expansion may be facilitated by the species exceptional tolerance to a wide range of abiotic stresses, including heavy metals, salinity, drought, antibiotics, and extreme temperatures, traits that likely enhance persistence in managed agricultural environments. In support of its ecological versatility, *P. azotoformans* has also been implicated in the blue discoloration of rabbit carcasses (Circella et al., 2022) and detected in food-processing environments (Thomassen et al., 2022). Overall, *P azotoformans* appears to be a highly adaptable generalist, capable of colonizing diverse hosts and ecological niches. Its prevalence in mushroom blotch may be facilitated by its ability to persist in acidic substrates (Mokrani et al., 2020; Wang et al., 2021), such as peat moss, a primary component of mushroom casing soil. However, their prevalence in bacterial blotch disease highlights the need of future studies on its evolutionary trajectory, host range, transmission pathways, and virulence mechanisms to understand its role in emerging disease complexes across agricultural and food systems. In addition to *P. azotoformans*, we also reported the first association of *P. pergaminensis* with mushroom blotch. This species was originally isolated from the wheat rhizosphere during tillering (Díaz et al., 2022) and is used as a bio-stimulant to enhance crop performance. While not previously linked to mushroom pathogenesis, *P. pergaminensis* possesses genomic traits consistent with rhizosphere competence, including phosphate and zinc solubilization and the secretion of extracellular lytic enzymes such as phospholipases and proteases (Díaz et al., 2022; Xiang et al., 2025) which may similarly facilitate colonization and symptom development on mushroom tissues.

The mechanisms underlying discoloration and pitting caused by *Pseudomonas* species remain poorly understood. However, accumulating evidence suggests that extracellular metabolites, such as cyclic lipopeptides (tolaasin), siderophores, degradative enzymes (proteases and lipases), and volatiles (tovsins, DMDS), play central roles in tissue degradation and corrosion processes (Lo Cantore et al., 2015; Soler-Rivas et al., 1999). These compounds mediate microbial competition, surface colonization, and host-pathogen interactions, all of which may contribute to blotch-like symptoms on mushroom caps. To explore this biochemical basis, we employed antiSMASH to predict the families of secondary metabolites encoded by all isolates and to assess their diversity and distribution. Heterogeneity in BGCs was evident across isolates, encompassing CLPs, siderophores, antibiotics, and volatile compounds. CLPs represent a major class of multifunctional secondary metabolites predominantly produced by *Pseudomonas* spp. Among the CLPs, WLIP has been identified as a diagnostic marker for the interaction between tolaasin and the cyclic WLIP produced by *Pseudomonas “reactans”* (Rokni-Zadeh et al., 2011), which manifests as a dense white line when the two strains are grown adjacently on agar. The white line reaction is highly structure-specific; WLIP analogues such as viscosin, containing L-Leu instead of D-Leu at position 5, and viscosinamide, a viscosin variant with D-Gln replacing D-Glu at position 2, fail to induce a white line in the presence of *P. tolaasii* (Rokni-Zadeh et al., 2011; Rokni-Zadeh et al., 2013), highlighting the strict molecular complementarity between WLIP and tolaasin. However, accumulating evidence indicates that this interaction is not exclusive. CLPs other than WLIP, including orfamides A, viscosin, and massetolide A, can also form visible white lines with tolaasin (Henkels et al. 2014). Similarly, sessilins, which are structurally related to tolaasin, induce white line formation through co-precipitation with orfamides (D’aes et al., 2014). Further evidence suggests that WLIP secretion is not restricted to *P. reactans*; *P. fluorescens* and *P. putida* strains also produce WLIP (Rokni-Zadeh et al., 2012). In the present study, we observed similar results related to WLIP. *Pseudomonas pergaminensis*, which produces viscosin and orfamides B but no detectable WLIP, formed a clear white line with *P. tolaasii* BP1106.

*P. azotoformans*, despite limited CLP production, also induced a white line. Notably, *P. aeruginosa* LMG 1272, which lacks canonical CLP biosynthetic genes associated with WLR, nonetheless forms a precipitate when co-cultured with *P. tolaasii* (Salari et al., 2020). Additionally, *Pseudomonas* species cluster 2 isolates (SM391, SM393, and SM396), which are positive for viscosin production, produced detectable white lines when confronted with *P. tolaasii* BP1106. These observations suggest that WLR formation is not strictly dependent on a single WLIP-tolaasin pair. Rather, variations in peptide stereochemistry, fatty acid chain length, or CLP production levels may influence diffusion and co-precipitation dynamics, resulting in differences in white line intensity and morphology. Siderophore BGCs, including EDHA, yersiniabactin, Ni-siderophore, enantio-pyochelin, and azotobactin D, were distributed among the isolates, consistent with the central role of iron acquisition in bacterial fitness (Page, 2013; Jimenez et al., 2010). However, the canonical siderophore pyoverdine, commonly associated with fluorescent *Pseudomonas* species, was not detected at sufficient similarity levels in any of the genomes. Furthermore, although tolaasin-like clusters were detected with moderate similarity (50–75%) in *P. pergaminensis*, *P. yamanorum*, and species cluster 3, white-line agar assays confirmed the absence of tolaasin activity. Taken together, these results highlight both the utility and the limitations of *in silico* BGC predictions: while they provide valuable chemotaxonomic signatures and candidate loci of interest, functional validation including domain-level annotation, proteomics, metabolomic analyses is necessary to confirm metabolite production and their direct contributions to virulence. Comparative pangenome analysis of *Pseudomonas tolaasii*, *P. pergaminensis*, *P*. “*gingeri*” and *P. azotoformans* revealed that only *P. azotoformans* possessed a significantly larger accessory genome relative to its core genome. This expanded accessory fraction corresponds with its broad ecological distribution, siderophore production, and environmental tolerance, reflecting high genomic plasticity for rapid adaptation. To assess the genetic diversity and expansion potential of the four *Pseudomonas* species, we analyzed their pan-genome trajectories using Tettelin et al.’s (2008) power-law regression model, based on Heaps’ Law. In this framework, the number of new genes discovered with each additional genome sequenced is described mathematically by *Yp = Ax^B + C*, where γ (exponent B) indicates pan-genome openness: γ > 0 denotes an open pan-genome, with genes continuing to accumulate, while γ < 0 denotes a closed pan-genome, where new gene discovery slows substantially. The Estimated γ values were 0.049 (*P. tolaasii*), 0.056 (*P. pergaminensis*), 0.046 (*P. “gingeri”*), and 0.085 (*P. azotoformans*). According to the power-law framework results indicate that all species possess theoretically open pangenomes, consistent with continued gene acquisition as additional genomes are incorporated. However, the relatively low γ values suggest that these pangenomes may be nearing closure. These interpretations should be approached with caution, as limited genome sampling can affect the reliability of power-law extrapolation. Consistent with our findings, previous studies have also reported an open pangenome for *Pseudomonas tolaasii* (Liu et al., 2022), although comprehensive comparative data across related species remain limited.

While the study is limited by a narrow temporal window and sampling restricted to two southern U.S. farms, future research will incorporate comparative analyses of both symptomatic and asymptomatic mushrooms, following the approach of Martins et al. (2020), to establish baseline microbiome diversity. The findings presented here enhance the current understanding of blotch disease etiology, revealing a greater taxonomic and functional diversity among the pathogens than previously recognized, and extending global insights into the complex bacterial community responsible for mushroom blotch. To our knowledge, this study provides the first evidence that *P. azotoformans*, *P. pergaminensis*, *P. monsensis*, *P. tensinigenes*, and *P. simiae* cause blotch in *Agaricus bisporus*, overturning the long-held assumption that the disease is primarily driven by *P. tolaasii* and *P. “gingeri”*. Species identities were robustly resolved through ANI, TYGS, GTDB-TK and GGDC. Secondary metabolite profiling and white-line assays further uncovered molecular features likely contributing to virulence and competitive fitness. The substantial diversity among pathogens, with *P. azotoformans* showing pronounced genomic expansion indicative of broad ecological adaptability. As newly emerging pathogens become increasingly prevalent, strain-level diagnostics and genomic surveillance will be critical for early detection and improved disease management in commercial mushroom production.

## 5. CREDIT AUTHORSHIP CONTRIBUTION STATEMENT

### Author Contributions

S.J.M. obtained the funding source; S.D.M., S.J.M., and R.G. designed the research; S.D.M. performed experiments; S.J.M. and M.L. assisted with experimental work; J.H. contributed to bioinformatics analyses; S.D.M. wrote the manuscript; R.G., and S.J.M reviewed the manuscript.

## 7. ACKNOWLEDGMENTS

The authors also acknowledge Dr. Fabiano Perina for the assistance with the rep-PCR assay and Dr. Carolee Bull who kindly shared the bacterial strains BP1304, BP1106 and *P. florescens* Pf-5 used in this study.

## 9. DATA AVAILABILITY

The whole genome data generated during the current study are available in the NCBI under Bio Project PRJNA1377199, with accession numbers SAMN53759319 to SAMN53759375.

---

**Supplementary Figure 1.**
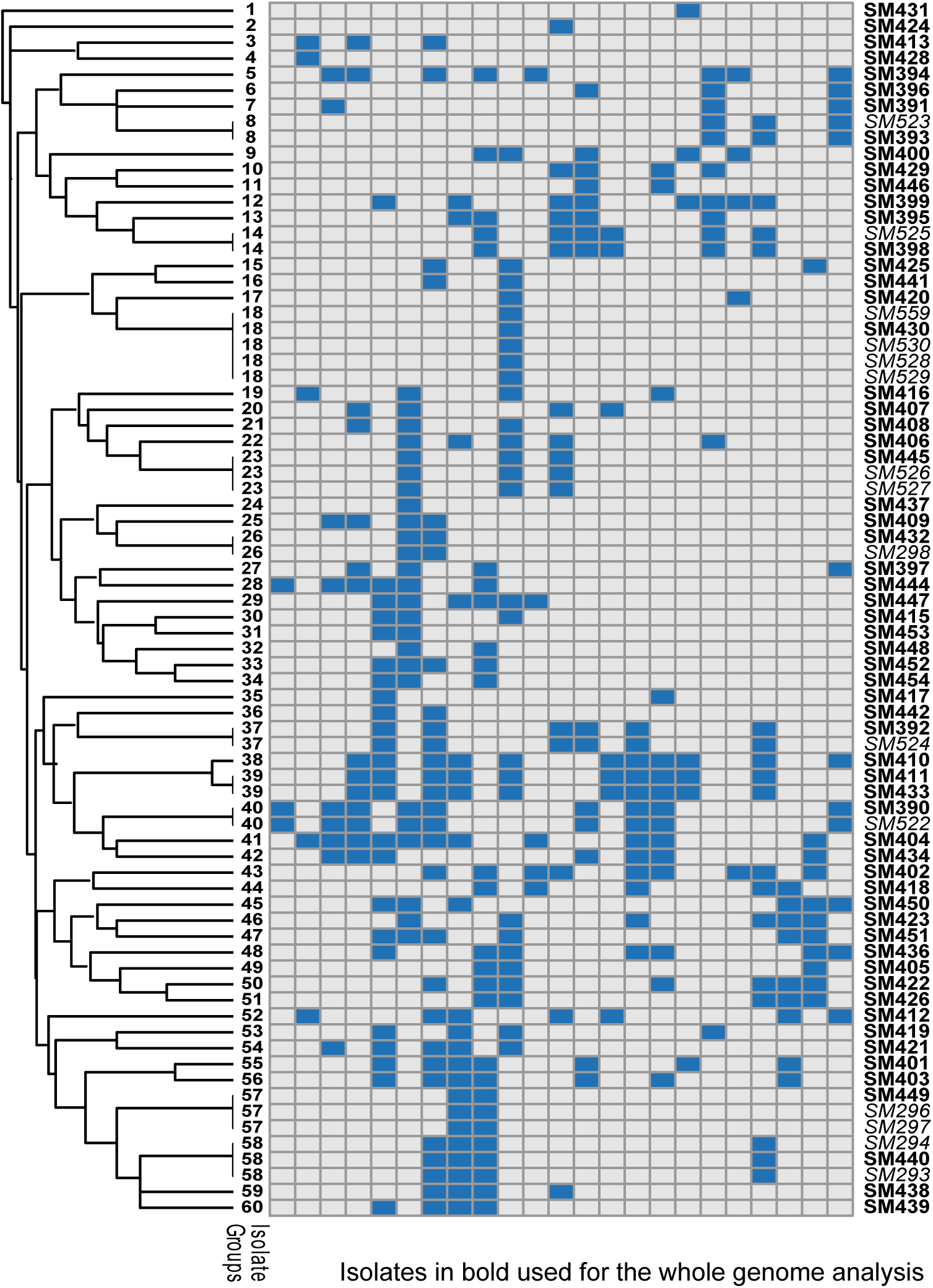
Genetic diversity of *Pseudomonas* isolates based on BOX PCR profiles. BOX-PCR fingerprinting of 77 bacterial isolates produced banding patterns on agarose gel. A binary data matrix, constructed from the presence or absence of 23 distinct BOX-PCR bands, was used for clustering analysis. The resulting dendrogram revealed 60 clusters at a 90% similarity threshold. Based on these DNA fingerprinting patterns, 15 strains with identical BOX-PCR profiles were isogenic and excluded from further analyses. Isolates shown in bold were selected for whole-genome sequencing.

